# Gene Set-Based Analysis of the Endosomal Sorting Processes Cargo Selection and Membrane Tubulation with Human Reward System Reactivity

**DOI:** 10.1101/2023.07.18.549434

**Authors:** Jens Treutlein, Bernd Krämer, Monika Rex-Haffner, Swapnil Awasthi, Stephan Ripke, Elisabeth B. Binder, Oliver Gruber

## Abstract

Dysfunction of the dopaminergic reward system has been implicated in the pathophysiology of several neuropsychiatric disorders. The endosomal network encompasses important processes related to neurotransmission in dopamine neurons, e.g, endocytosis, sorting, recycling and degradation of receptors. In the present study, we investigated whether genetic variation influencing the endosomal sorting machinery, in particular cargo selection and membrane tubulation, may impact on the activation strength in key regions of the mesolimbic reward system, i.e. the ventral tegmental area (VTA) and the nucleus accumbens (NAc).

To test our hypothesis, the VTA and NAc responses to conditioned reward stimuli were investigated using the ‘desire-reason dilemma’ (DRD) paradigm during functional magnetic resonance imaging (fMRI). Association of these neural responses with a set of genetic variants related to endosomal sorting processes were tested in two independent samples (N = 182; N = 214).

In the first sample, the gene set was associated with both VTA and NAc responses to conditioned reward stimuli [empirical P -values: R-VTA 0.0036; R-NAc 0.0016; L-NAc 0.0094], while the effect in the R-VTA could be replicated in the second sample [empirical P -value: R-VTA 0.0443] at the level of the gene set. For the NAc, an additional exploratory analysis of a patient-only subsample of the first sample (N = 64) suggested that the gene set may express its effect in this brain region predominantly in patients.

These findings provide first evidence that the endosomal sorting processes cargo selection and membrane tubulation influence neural responses of the reward system to conditioned stimuli. Further studies are required to clarify the role of endosomal sorting processes in the pathophysiology of neuropsychiatric disorders.

## INTRODUCTION

Dysfunction of the reward system is well known to play a prominent role in psychiatric disorders. The core of the mesolimbic dopamine pathway is represented by a population of dopamine-containing neurons in the ventral tegmental area (VTA) that project to the nucleus accumbens (NAc) (Dichter, Damiano, & Allen, 2012; Russo & Nestler, 2013). In humans, clear evidence for disturbances in these regions for psychiatric disorders were provided by studies of reward-specific paradigms in functional magnetic resonance imaging (fMRI) studies (Deserno, Schlagenhauf, & Heinz, 2016; Goya-Maldonado et al., 2016; Qi et al., 2021; Richter et al., 2015; Trost et al., 2014).

In the reward system, as in tissues in general, protein homeostasis is essential for cell function. In this context, the multi-component protein complexes of the endosome function as a protein-sorting hub. After surface molecules have been internalized from the cell surface, they traffic through an endosomal series of tubulovesicular compartments and are either sorted for recycling back to the plasma membrane, to the trans-Golgi-network, or are targeted to lysosomal degradation. Under physiological conditions, a balanced coordination exists between the sorting pathways; however, dysregulation of one or more of these pathways results in aberrant protein sorting which may lead to cell toxicity (Vagnozzi & Pratico, 2019).

In neurons, endosomes fulfill specific functions that are related to neurotransmission. At the presynapse, the endosomal network regulates the pool of synaptic vesicles that can be released, by either generating new vesicles or by degrading vesicles that are not needed or that are damaged. At the postsynapse, the endosomal network regulates receptor recycling and endocytosis, with the number of receptors on the post synaptic membrane influencing the strength of the signal being propagated (Plooster, Brennwald, & Gupton, 2022; Schmidt & Haucke, 2007; Vieira, Rito, Correia-Neves, & Sousa, 2021). Thus, via endosomal trafficking, neurons can regulate the reuptake of neurotransmitters and signaling from neurotransmitter receptors, and alterations in this process results in changes of neuronal function and plasticity.

Previous research has functionally connected the endosomal trafficking process with dopaminergic neurotransmission, due to an important role of this process for regulation of key proteins of dopamine-releasing neurons (e.g. dopamine transporter, (Block et al., 2015)) and their postsynaptic neurons (e.g. dopamine receptors, (Bartlett et al., 2005; Dumartin et al., 2000; Li et al., 2012; Zhang et al., 2012)). Furthermore, modification of endosomal protein Vps35 disturbed dopamine function (Ishizu et al., 2016). Additionally, endosomal sorting has been recognized to play a role for the etiology of neuropsychiatric disorders that are related to dopaminergic dysfunction, e.g., schizophrenia (Plooster et al., 2022) and Parkinson’s disease (Cunningham & Moore, 2020; Vagnozzi & Pratico, 2019).

In the highly complex endosomal sorting process, one specific aspect is the selection and partitioning of cargo proteins into tubular transport intermediates, which requires the interaction of two multi-protein complexes that collect and package the cargo, the ’retromer’ complex and the ’Wiskott– Aldrich syndrome protein and SCAR homologue’ (WASH) complex.

A number of genes whose products play a role in the selection and partitioning of cargo proteins in the endosomal sorting machinery are known (Freeman, Hesketh, & Seaman, 2014). Testing for the aggregated contribution of genetic variants in groups of genes is an approach to circumvent the extreme correction for multiple testing in single-marker analysis, under the assumption that functionally related genes contain multiple variants that disrupt function. An example for this are gene set-based methods (Wang, Li, & Hakonarson, 2010), typically in the form of consecutive enzymes in a biochemical reaction or in the form of functional groups, by grouping genes whose protein products interact. Such a group can be profoundly investigated by polymorphisms that are located within the coding sequence of the genes, i.e. non-synonymous variants that exchange the amino acid (or may influence splicing) (Tabor, Risch, & Myers, 2002), and synonymous variants that play multifunctional structural and regulatory roles by defining secondary structure and stability of the mRNA, influence splicing, translation rate, post-translational modification, or polypeptide chain folding (Shabalina, Spiridonov, & Kashina, 2013).

We hypothesized that individuals carrying genetic variants whose products influence cargo selection and membrane tubulation, display changes in subcortical reward processing in humans. For this purpose, we investigated effects of these genetic variants in two independent samples using fMRI and genetic methods. We conducted gene set-based tests to investigate if the joint activity of these genetic variants influence the activation strength of brain regions within the reward system in humans.

## MATERIALS & METHODS

### Subjects and Genotyping

The two samples that were used in our study contained subjects that performed the same experimental procedure, and the same fMRI data acquisition and phenotype extraction was applied to both. Sample 1 (N = 182) consisted of adult patients with psychiatric disorders that were recruited in the context of clinical studies, and adult unaffected individuals that were recruited as controls for these studies (Goya-Maldonado et al., 2016; Richter et al., 2015; Trost et al., 2014). Additional exploratory analyses were conducted in patients (N = 64), as a patient-only subsample of sample 1. Sample 2 (N=214) consisted of a collection of healthy adult young individuals, as previously described (Trost et al., 2016). All participants were of European ancestry. Every subject provided written informed consent. The study was carried out in accordance with the Declaration of Helsinki and was approved by the local ethics committee.

Saliva was collected using Oragene DNA devices (DNA Genotek, Ottawa, Ontario, Canada), and DNA was isolated with standardized protocols. Genotyping was performed using Illumina Genotyping BeadChips (OmniExpress). This was followed by genotype imputation using EAGLE/MINIMAC3 with Haplotype Reference Consortium (HRC) data, and by quality control (exclusion of cryptically related pairs; exclusion of population outliers). These procedures were consistently applied to the individuals of both samples.

### Gene Set-Based Association Analysis

Gene set-based association analysis jointly tests genetic variants as a single gene set rather than as single markers, to reduce the number of required corrections due to multiple testing. For analysis, genetic variants were selected that were broadly supported as being located in the coding sequence of retromer and WASH genes in public databases (Bhagwat, 2010; Raudvere et al., 2019; UniProt, 2023).

- rs3762672 [T/G, 3:132218623 hg19] nonsynonymous variant [A(GCT)->S(TCT)] in the *DNAJC13* gene [alias Receptor Mediated Endocytosis 8; RME-8] on chr3q22.1.
- rs17157971 [T/C, 10:46224381 hg19] synonymous variant [D(GAC)->D(GAT)] in the *WASHC2C* gene [alias Family With Sequence Similarity 21 Member C; FAM21C] on chr10q11.22.
- rs3751129 [A/G, 12:102455729 hg19] synonymous variant [(D(GAC)->D(GAT)] in the *WASHC3* gene [alias Coiled-Coil Domain-Containing protein 53; CCDC53] on chr12q23.2.
- rs1663564 [G/A, 12:105546172 hg19] nonsynonymous variant [V(GTA)->I(ATA)] in the *WASHC4* gene [alias KIAA1033; alias Strumpellin And WASH-Interacting Protein; SWIP] on chr12q23.3.
- rs12432539 [C/T, 14:35062312 hg19] synonymous variant [E(GAG)->E(GAA)] in the *SNX6* gene on chr14q13.1.
- rs1802376 [A/G, 15:64428559 hg19] nonsynonymous variant [D(GAC)->N(AAC)] in the *SNX1* gene on chr15q22.31.

In the analysis method used for the present study, the PLINK set-test, first a single variant analysis in the original set is performed. From the statistics of the single variants contained in the set, a mean statistic is calculated. The same analysis is then performed with a certain number of simulated sets with a permuted phenotype status of the individuals. From these analyses, an empirical P -value (‘EMP1’) is retrieved by calculation of the number of times that the test statistic of the simulated sets exceeds the test statistic of the original set (Deelen et al., 2013). PLINK version v1.90b6.9 (Chang et al., 2015) was used with the adaptive procedure (flag ‘--assoc perm set-test’), without exclusion of any of the six variants due to P -value or linkage disequilibrium thresholds.

### Desire-Reason-Dilemma Paradigm

All subjects had performed the ‘desire-reason dilemma’ (DRD) paradigm allowing systematic investigation of systems-level mechanisms of reward processing in humans. In this fMRI experiment, participants initially undergo an operant conditioning task. Following this task, subjects perform the DRD paradigm. For the present study, one part of the DRD paradigm was used that assesses the bottom-up activation of the reward system by conditioned stimuli. For more details on the paradigm see Diekhof & Gruber, 2010; Diekhof et al., 2012 (Diekhof & Gruber, 2010; Diekhof et al., 2012).

### fMRI Data Acquisition

fMRI was performed on a 3-Tesla Magnetom TIM Trio Siemens scanner (Siemens Healthcare, Erlangen, Germany) equipped with a standard eight-channel phased-array head coil. First, a T1-weighted anatomical data set with 1 mm isotropic resolution was acquired. Parallel to the anterior commissure– posterior commissure line, thirty-one axial slices were acquired in ascending direction for fMRI (slice thickness = 3 mm; interslice gap = 0.6 mm) using a gradient-echo echo-planar imaging sequence (echo time 33 ms, flip angle 70°; field-of-view 192 mm, interscan repetition time 1900 ms).

In 2 functional runs, 185 volumes each were acquired. Subjects responded via button presses on a fiber optic computer response device (Current Designs, Philadelphia, Pennsylvania, USA), and stimuli were viewed through goggles (Resonance Technology, Northridge, California, USA). Presentation Software (Neurobehavioral Systems, Albany, California, USA) was used to present the stimuli in the scanner. Functional images were preprocessed and analyzed with SPM12 (https://www.fil.ion.ucl.ac.uk/spm/) using a general linear model. The study design was event-related and only correctly answered trials were included in the analysis. Linear t-contrasts were defined to assess activation effects elicited by the conditioned reward stimuli.

### fMRI Phenotype Extraction

For the subsequent genetic association analyses, neuroimaging phenotypes were determined. The individual mean beta estimates from the *a priori* regions of interest (ROIs), namely the VTA and the NAc, were extracted with SPM12. Beta extraction for each participant was performed using predefined MNI coordinates of both brain regions emerging from previous publications: ±9 -21 -15 for the bilateral VTA and ±12 12 -3 for the bilateral NAc (Diekhof et al., 2012; Jakob, 2012; Trost et al., 2016). To account for interindividual functional neuroanatomical variation that may be insufficiently covered by spatial normalization processes in the standard preprocessing pipeline, a box with a 3 × 3 × 3 mm^3^ dimension around the *a priori* MNI coordinates was applied to determine the individual maximum of reward-related brain activation within each subject.

## RESULTS

Genotypes of the variants could be determined for n=182 subjects in sample 1, and for n=214 subjects in sample 2, and were in Hardy-Weinberg Equilibrium (HWE p>0.05). Allele frequencies in our samples were in line with those of the 1000 genomes project phase 3 CEU population (1KGP-3-CEU; http://grch37.ensembl.org/Homo_sapiens/Info/Index), which demonstrates that genotype determination was reliable in our samples (Table S1, Table S2).

### Gene Set-Based Association Analysis

Gene set-based analyses were used to investigate the joint contribution of coding sequence variants in the endosomal sorting processes cargo selection and membrane tubulation to the extent of activation of the reward system. For all regions of interest in sample 1, we observed a significant joint effect of the variants in the gene set (EMP1_R-VTA_ = 0.0036; EMP1_L-VTA_ = 0.0451; EMP1_R-NAc_ = 0.0016; EMP1_L-NAc_ = 0.0094). Of these brain regions, three (R-VTA, R-NAc, L-NAc) survived correction for four comparisons at p<0.05. In addition, the effect in the R-VTA could be replicated in sample 2 (EMP1_R-VTA_ = 0.0443). Full gene set test results for both brain regions and both samples are shown in Table 1.

**Table 1.** Set-based and single marker-based association results in samples 1 and 2. For set-based analyses EMP1 -values (set-test empirical P – values) are shown; for single marker analyses effect alleles (EA) and single marker P – values (P) with effect sizes (BETA) in parentheses are shown; EMP1-values and single marker P -values < 0.05 in bold.

| *Sample 1* |  |  |  |  |  |
| --- | --- | --- | --- | --- | --- |
|  |  | EMP1<sub>R-VTA</sub>: **0.0036** | EMP1<sub>L-VTA</sub>: **0.0451** | EMP1<sub>R-NAC</sub>: **0.0016** | EMP1<sub>L-NAC</sub>: **0.0094** |
| *Variant ID* | *EA* | *P<sub>R-VTA</sub> (BETA<sub>R-VTA</sub>)* | *P<sub>L-VTA</sub> (BETA<sub>L-VTA</sub>)* | *P<sub>R-NAC</sub> (BETA<sub>R-NAC</sub>)* | *P<sub>L-NAC</sub> (BETA<sub>L-NAC</sub>)* |
| *rs3762672* | T | **0.0188** (-0.4675) | 0.1918 (-0.2531) | 0.7540 (-0.0939) | 0.7760 (-0.0807) |
| *rs17157971* | T | 0.6085 (-0.2721) | 0.0747 (-0.9159) | 0.4448 (-0.6051) | 0.5819 (-0.4130) |
| *rs3751129* | A | **0.0147** (-0.5864) | **0.0059** (-0.6499) | **0.0383** (-0.7574) | 0.0661 (-0.6328) |
| *rs1663564* | G | 0.1496 (0.5277) | 0.5671 (0.2041) | 0.1401 (0.8057) | 0.2105 (0.6480) |
| *rs12432539* | C | **0.0155** (-0.5673) | 0.1413 (-0.3363) | **0.0052** (-0.9745) | **0.0139** (-0.8120) |
| *rs1802376* | A | 0.6639 (0.3001) | 0.6775 (0.2789) | **0.0028** (3.0450) | **0.0099** (2.4930) |
| *Sample 2* |  |  |  |  |  |
|  |  | EMP1<sub>R-VTA</sub>: **0.0443** | EMP1<sub>L-VTA</sub>: 0.0897 | EMP1<sub>R-NAC</sub>: 0.3846 | EMP1<sub>L-NAC</sub>: 0.1443 |
| *Variant ID* | *EA* | *P<sub>R-VTA</sub> (BETA<sub>R-VTA</sub>)* | *P<sub>L-VTA</sub> (BETA<sub>L-VTA</sub>)* | *P<sub>R-NAC</sub> (BETA<sub>R-NAC</sub>)* | *P<sub>L-NAC</sub> (BETA<sub>L-NAC</sub>)* |
| *rs3762672* | T | **0.0007** (0.6239) | **0.0195** (0.3882) | 0.3121 (0.1992) | 0.1334 (0.2933) |
| *rs17157971* | T | 0.6217 (-0.2371) | 0.9300 (0.0382) | 0.5958 (0.2703) | 0.5808 (0.2795) |
| *rs3751129* | A | 0.6136 (-0.1172) | 0.1462 (-0.3042) | 0.0750 (0.4367) | 0.4515 (-0.1840) |
| *rs1663564* | G | 0.6006 (-0.1529) | 0.5422 (-0.1609) | 0.2286 (-0.3725) | **0.0118** (-0.7694) |
| *rs12432539* | C | 0.9615 (0.0115) | 0.6721 (-0.0913) | 0.1628 (-0.3526) | 0.3414 (-0.2390) |
| *rs1802376* | A | 0.2498 (-0.9885) | 0.1132 (-1.2280) | 0.4188 (-0.7374) | 0.7479 (-0.2914) |

### Single-Variant Association Analysis

In addition to computation of an empirical P -value of a gene set, PLINK 1.9 conducts association analyses of single markers contained in the set. In the three gene set tests that survived testing for multiple comparisons (R-VTA, R-NAc and L-NAc regions in sample 1), four genetic variants reached significance at P < 0.05: rs3762672 (P = 0.0188), rs3751129 (P = 0.0147) and rs12432539 (P = 0.0155) were associated with the strength of activation in the R-VTA region, whereas rs3751129 (P = 0.0059) influenced the strength of response in the L-VTA. rs3751129 (P = 0.0383), rs12432539 (P = 0.0052) and rs1802376 (P = 0.0028) were associated with the response in the R-NAc region, and rs12432539 (P = 0.0139) and rs1802376 (P = 0.0099) influenced L-NAc reactivity. Because rs3751129 and rs12432539 were involved in the association of two associated brain regions, R-VTA and R-NAc, their respective genes *WASHC3* and *SNX6* constitute the strongest findings of our analysis. Consistent across R-VTA and R-NAc brain regions, the risk allele of rs3751129 for increased activation was the G-allele, and the risk allele of rs12432539 for increased activation was the T-allele. Beside these two genes, *DNAJC13* variant rs3762672 showed nominal significance across both samples (R-VTA region in sample 1: P = 0.0188; R-VTA in sample 2: P = 0.0007), but with a reversal of the allele. Allele reversals sometimes occur in genetic association studies and have been described in detail with possible reasons supporting their occurrence (Gruber, Genissel, Macdonald, & Long, 2007; Lin, Vance, Pericak-Vance, & Martin, 2007; Maher, Reimers, Riley, & Kendler, 2010; Ober, 2016; Zaykin & Shibata, 2008). Full single marker association test results for both brain regions and both samples are given in Table 1.

### Exploratory Set-Based and Single Marker-Based Association Analyses in the Patient-Only Subsample

Results retrieved from sample 1 (which consisted of both patients and controls) could be replicated in the healthy individuals of sample 2 only for the R-VTA. A possible explanation is that our gene set contains largely pathogenic factors, which lead to changes of reward system reactivity predominantly in patients. Patients are exposed to more environmental risk factors compared to healthy subjects, and one could assume that these risk factors may interact with the genetic variants contained in the gene set leading to increased effect size. Although reward system reactivity is a polygenic trait, the situation may be reminiscent of monogenic phenylketonuria, where uptake of phenylalanine as an environmental factor interacts with the mutant phenylalanine hydroxylase gene genotype to result in impaired postnatal cognitive development only in patients (Amberger, Bocchini, Scott, & Hamosh, 2019).

Therefore, in order to further explore the effects of our gene set in patients, we compiled a subsample consisting of the 64 patients contained in sample 1. For this pure patient sample, set-based results remained significant for both NAc regions (while the corresponding effect on the VTA could not be confirmed). In fact, the observed effects of variant rs1802376 in single-marker analysis improved when focusing on patients only (full sample 1 in R-NAc: P = 2.8e-03, patient-only subsample of sample 1 in R-NAc: P = 6.111e-05; full sample 1 in L-NAc: P = 9.9e-03, patient-only subsample of sample 1 in L-NAc: P = 4.159e-05) (Table S3).

The findings of this exploratory analysis suggests that in the NAc the pathophysiological effects to which the gene set contributes may be predominantly expressed in patients.

## DISCUSSION

The goal of our study was to test the hypothesis that allele-related variation in genes of the endosomal sorting processes cargo selection and membrane tubulation is associated with the reactivity of subcortical key regions of the mesolimbic reward system in response to reward stimuli. We used two well-suited samples to address the question how genetic variation affects the functional characteristics of the brain reward system. The specific subprocesses of endosomal sorting that were tested in the present study involve two large protein complexes, the retromer complex and the WASH complex. Using a set-based test, we detected significant effects of relevant genes of these complexes that contribute to the endosomal sorting subprocesses in both analysed regions of the reward system.

SNX1 and SNX6 of our gene set are components of retromer, which is a heteropentameric complex in humans, consisting of sorting nexin (SNX) and vacuolar sorting associated proteins (VPS). The SNX dimer plays a role in deformation of the endosomal membrane and generates the tubules for cargo transport. The other retromer component, the cargo-selective complex (CSC) is composed of a trimer consisting of three vacuolar sorting associated proteins, and transiently associates with the SNX dimer.

WASHC2C, WASHC3 and WASHC4 in our gene set were aquired from WASH. WASH is a pentameric complex that generates a branched actin network stabilizing an endosomal microdomain which serves to concentrate specific cargoes. The activity between the retromer and WASH complexes is coordinated by DNAJC13, a protein that binds to SNX1 of the retromer complex and to a WASH complex subunit (Freeman et al., 2014).

Of the endosomal sorting components tested in the present study, the most robust signals for increased responses of the reward system were retrieved from *WASHC3* and *SNX6*, because these two genes were implicated in the modulation of brain activity in both key regions, i.e. the VTA and the NAc. Additonally, the association of *DNAJC13* is of particular interest because this gene was involved in R-VTA reactivity across both samples. Furthermore, *SNX1* emerged from the patient-only analysis with increased signal strength. Gene products of all these most consistently associated genes of our study play specific roles in endosomal sorting and tubulation processes, and three of them (*WASHC3, DNAJC13, SNX1*) were in the past plausibly linked to traits related to dopaminergic dysfunction:

The first of our genes, namely the *WASHC3* gene which was associated with activation of VTA and NAc regions in our single marker tests, was suggested to function as a nucleation-promoting factor at the surface of endosomes where it was suggested to induce actin polymerization (https://www.uniprot.org/uniprotkb/Q9Y3C0/entry#function). The altered activation of the VTA in the present study may at least in part be explained by the reported influence of rs3751129 on regulation of the volume of the midbrain (p=1.83e-09) (Elvsashagen et al., 2020).

The second gene, *DNAJC13*, which is the only gene implicated across both of our samples, is of particular importance for coordination between retromer and WASH, because DNAJC13 regulates the packaging process spatially and temporally, by ensuring that the requisite elements of the transport machinery are present at the microdomain (Freeman et al., 2014; Fujibayashi et al., 2008; Girard, Poupon, Blondeau, & McPherson, 2005). rs3762672 mediates an amino-acid change of the hydrophobic residue alanine to the polar residue serine at amino acid position 1463 in the human protein, and has been observed in Parkinson’s disease with dementia in a single case, however with unclear relevance (Smaili et al., 2020). The involvement of DNAJC13 in dopaminergic neurotransmission in the present study may provide a mechanistic explanation of previous reports for *DNAJC13*, where rare missense variants of this gene contributed to Parkinson’s disease (https://www.omim.org/entry/616361) (Yoshida et al., 2018), and gene expression and rare disruptive mutations were implicated in schizophrenia (Piras et al., 2019; Wu, Yao, & Luo, 2017).

The third gene, SNX1, which is a direct molecular interaction partner of DNAJC13, was associated with reactivity of both NAc regions, and showed a particularly evident association signal in the patient-only subsample. It therefore may point to a pathological condition, which however must be further investigated and replicated in future studies. rs1802376 mediates an amino-acid change of the negatively charged residue aspartic acid to the uncharged residue asparagine at amino acid position 466 in the human protein. This missense variant has been predicted as probably damaging for the SNX1 protein by PolyPhen-2 (Adzhubei et al., 2010). This provides an explanation for the association signal for this variant detected in the in the data of the present study, and in a mega-analysis of the Psychiatric Genomics Consortium Schizophrenia Working Group (p = 0.001837) (Schizophrenia Working Group of the Psychiatric Genomics, 2014).

The last gene, SNX6, influenced the activation of VTA and NAc regions in our single-marker analysis. Its gene product (together with its paralog SNX5) is known to form a banana-like structure that senses and generates membrane curvature and contacts dynactin, an activator of the microtubule motor dynein (Wassmer et al., 2009). Furthermore, the dimer recognizes a bipartite sorting signal (Hong, 2019). rs12432539 may influence alternative splicing of the *SNX6* messenger RNA in an allele-specific manner (Amoah et al., 2021; Nembaware et al., 2008), and the resulting qualitative alteration of the *SNX6* transcript may explain the association signal observed in the present study. In contrast to the other three endosomal sorting components implicated by our analyses, there are – to the best of our knowledge -no reports connecting SNX6 with dopamine. Therefore, our results may point to a previously undiscovered relevance of this sorting nexin in processes related to this neurotransmitter.

From the molecular systems perspective, the detected associations suggest that both macromolecular complexes, retromer and WASH, as well as the coordination of the activity between the two complexes, contribute to modulation of differential reactivity of the reward system. Genetic variation in endosomal sorting pathways may increase the vulnerability for neuropsychiatric disorders via activity changes of the mesolimbic dopamine system. Thus, our gene set-based tests of endosomal sorting components could be followed-up for a possible role in the pathogenesis of more specific forms of schizophrenia and/or affective disorders that are related to reward system dysfunction. If the gene set shows increased effects in those subforms in which dopaminergic mechanisms play a major role, this would strongly corroborate endosomal sorting as an underlying pathomechanism contributing to these disorders.

Another particularly interesting aspect for future investigations are cellular processes that are tightly connected to cargo selection and membrane tubulation, for example the generation of synaptic vesicles that are budding from the endosome (Gorenberg & Chandra, 2017; Limanaqi, Biagioni, Gambardella, Ryskalin, & Fornai, 2018), or the initiation of autophagy from the endosome (Besemer et al., 2021; Limanaqi et al., 2018). If these functionally connected processes also display association signals, it would indicate that a broader array of molecular pathomechanisms related to endosomal trafficking influence psychiatric disorders via reward processing.

In sum, the present study is the first fMRI study in humans that investigated the endosomal sorting processes cargo selection and membrane tubulation in the context of reward system reactivity. Our findings indicate that these processes influence neural responses to conditioned reward stimuli in the brain regions VTA and NAc.

## Acknowledgements

The authors gratefully acknowledge the data storage service SDS@hd supported by the Ministry of Science, Research and the Arts Baden-Württemberg (MWK) and the German Research Foundation (DFG) through grant INST 35/1503-1 FUGG. The authors further acknowledge support by the High Performance and Cloud Computing Group at the Zentrum für Datenverarbeitung of the University of Tübingen, the state of Baden-Württemberg through bwHPC and the German Research Foundation (DFG) through grant no INST 37/935-1 FUGG. We thank all subjects who participated in this study. We also thank Maria Keil, Center for Translational Research in Systems Neuroscience and Psychiatry, Department of Psychiatry and Psychotherapy, Georg-August-University Göttingen, Göttingen, for support in recruitment and fMRI investigation of volunteers.

## Compliance with ethical standards

The study was carried out in accordance with the Declaration of Helsinki and was approved by the local ethics committees, of the Medical Faculty of Göttingen University (number 14/3/09, date 02.07.2009) and of the Medical Faculty of Heidelberg University (number S-123/2016, date 09.03.2016). All participants provided written informed consent.

## SUPPLEMENTARY INFORMATION

Supplementary information accompanies the paper.

