## Supplementary Tables for "Gene Set-Based Analysis of the Endosomal Sorting Processes Cargo Selection and Membrane Tubulation with Human Reward System Reactivity"

[1]Section for Experimental Psychopathology and Neuroimaging, Department of General Psychiatry, University Hospital Heidelberg, Germany; [2]Department Genes and Environment, Max-Planck-Institute for Psychiatry, Munich, Germany; [3]Department of Psychiatry and Psychotherapy, Charité - Universitätsmedizin, Berlin, Germany; [4]Stanley Center for Psychiatric Research, Broad Institute, Cambridge, MA, USA; [5]Analytic and Translational Genetics Unit, Massachusetts General Hospital, Boston, MA, USA.

#correspondence should be send to Professor Dr. Oliver Gruber;, Address: Voßstraße 4, 69115 Heidelberg, Germany; Phone number +49-6221-567511, Fax number +49-6221-566749.
ORCID: Jens Treutlein http://orcid.org/0000-0003-3952-8905

Supplementary Table 1. Genotype-related information in samples 1 and 2. Hom/Het/Hom: genotype counts; FREQ: minor allele frequency; HWE: Hardy-Weinberg equilibrium P - value; 1KGP-3: 1000 Genomes Project Phase 3 allele frequencies in the subsample of Utah residents (CEPH) with Northern and Western European ancestry (CEU).

*Sample 1*

| *Variant ID* | *Hom* | *Het* | *Hom* | *FREQ* | *HWE* | *1KGP-3* |
|---|---|---|---|---|---|---|
| *rs3762672* | GG:49 | GT:87 | TT:43 | T:0.483 | 0.765 | T:0.490 |
| *rs17157971* | CC:168 | CT:14 | TT: 0 | T:0.038 | 1.000 | T:0.045 |
| *rs3751129* | GG:109 | GA: 62 | AA: 8 | A:0.218 | 1.000 | A:0.217 |
| *rs1663564* | AA: 153 | AG: 28 | GG: 1 | G:0.082 | 1.000 | G: 0.071 |
| *rs12432539* | TT: 118 | TC: 52 | CC: 11 | C:0.204 | 0.113 | C:0.177 |
| *rs1802376* | GG: 174 | GA:8 | AA: 0 | A:0.022 | 1.000 | A:0.030 |

*Sample 2*

| *Variant ID* | *Hom1* | *Het* | *Hom2* | *FREQ* | *HWE* | *1KGP-3* |
|---|---|---|---|---|---|---|
| *rs3762672* | GG:54 | GT:108 | TT:51 | T:0.493 | 0.891 | T:0.490 |
| *rs17157971* | CC:197 | CT:17 | TT:0 | T:0.040 | 1.000 | T:0.045 |
| *rs3751129* | GG:123 | GA:84 | AA:7 | A:0.229 | 0.123 | A:0.217 |
| *rs1663564* | AA:175 | AG:35 | GG:4 | G:0.100 | 0.240 | G: 0.071 |
| *rs12432539* | TT:151 | TC:55 | CC:8 | C:0.166 | 0.320 | C:0.177 |
| *rs1802376* | GG:209 | GA:5 | AA:0 | A:0.012 | 1.000 | A:0.030 |

Supplementary Table 2. Genotype-related information in the patient-only subsample of sample 1. Hom/Het/Hom: genotype counts; FREQ: minor allele frequency; HWE: Hardy-Weinberg equilibrium P - value; 1KGP-3: 1000 Genomes Project Phase 3 allele frequencies in the subsample of Utah residents (CEPH) with Northern and Western European ancestry (CEU).

| *Patient-Only Subsample of Sample 1* | | | | | | |
|---|---|---|---|---|---|---|
| *Variant ID* | *Hom* | *Het* | *Hom* | *FREQ* | *HWE* | *1KGP-3* |
| *rs3762672* | GG:17 | GT:33 | TT:14 | T: 0.477 | 1.000 | T:0.490 |
| *rs17157971* | CC:58 | CT:6 | TT:0 | T: 0.047 | 1.000 | T:0.045 |
| *rs3751129* | GG:41 | GA:17 | AA:5 | A: 0.214 | 0.131 | A:0.217 |
| *rs1663564* | AA:49 | AG:14 | GG:1 | G: 0.125 | 1.000 | G: 0.071 |
| *rs12432539* | TT:43 | TC:18 | CC:3 | C: 0.188 | 0.678 | C:0.177 |
| *rs1802376* | GG:61 | GA:3 | AA:0 | A: 0.023 | 1.000 | A:0.030 |

Supplementary Table 3. Gene set-based and single marker-based association results in the patient-only subsample of sample 1. For set-based analyses EMP1 - values (set-test empirical P – values) are shown; for single marker analyses effect alleles (EA) and single marker P – values (P) with effect sizes (BETA) in parentheses are shown; EMP1-values and single marker P - values < 0.05 in bold.

| *Patient-Only Subsample of Sample 1* | | | | | |
|---|---|---|---|---|---|
| | | $EMP1_{R\text{-}VTA}$: 0.2353 | $EMP1_{L\text{-}VTA}$: 0.6667 | $EMP1_{R\text{-}NAc}$: **0.0031** | $EMP1_{L\text{-}NAc}$: **0.0020** |
| *Variant ID* | *EA* | $P_{R\text{-}VTA}$ ($BETA_{R\text{-}VTA}$) | $P_{L\text{-}VTA}$ ($BETA_{L\text{-}VTA}$) | $P_{R\text{-}NAc}$ ($BETA_{R\text{-}NAc}$) | $P_{L\text{-}NAc}$ ($BETA_{L\text{-}NAc}$) |
| *rs3762672* | T | **0.0289** (-0.7174) | 0.8385 (0.0606) | 0.5723 (-0.3448) | 0.3088 (-0.4957) |
| *rs17157971* | T | 0.7889 (-0.2134) | 0.3334 (-0.6820) | 0.6033 (-0.7559) | 0.6053 (-0.6016) |
| *rs3751129* | A | 0.1740 (-0.5012) | 0.0665 (-0.5945) | **0.0401** (-1.3670) | **0.0279** (-1.1630) |
| *rs1663564* | G | 0.4551 (0.3704) | 0.4071 (0.3646) | 0.3137 (0.9108) | 0.2321 (0.8628) |
| *rs12432539* | C | 0.9884 (-0.0059) | 0.4840 (0.2516) | **0.0441** (-1.4680) | 0.2390 (-0.6943) |
| *rs1802376* | A | 0.8511 (0.2064) | 0.6712 (0.4140) | **6.111e-05** (7.5520) | **4.159e-05** (6.1600) |
